# Genetic Diversity and Population Structure of Maize Doubled Haploid Lines from Drought and Low Nitrogen Tolerant Populations

**DOI:** 10.64898/2026.08.06.743182

**Authors:** Georgina Lala Ehemba, Beatrice E. Ifie, Bishwanath Das, Pearl Abu, Emmanuel Amponsah Adjei, Mathieu A.T. Ayenan, Ana Luisa Garcia-Oliveira, Priscilla Francisco Ribeiro, Manilal William, Pangirayi B. Tongoona, Eric Yirenkyi Danquah

## Abstract

Understanding the genetic diversity and population structure of breeding materials is essential for developing stress-resilient cultivars. In tropical maize, where drought and low soil nitrogen (low N) severely limit productivity, continuous development of tolerant varieties remains a priority. This study assessed the genetic diversity and population structure of 250 doubled haploid lines (DHLs) derived from five drought- and low N-tolerant tropical populations. Genotyping was performed using mid-density DArTseq markers, yielding 3,305 high-quality SNPs for analysis. Results revealed a moderate level of diversity among the DHLs, with an average genetic distance of 0.39, a polymorphism information content (PIC) of 0.33, and a minor allele frequency (MAF) of 0.29. These values reflect substantial allelic variation, important for identifying complementary parental combinations in hybrid development. Discriminant analysis of principal components (DAPC) grouped the DHLs into five distinct clusters, largely corresponding to their source populations, although some admixture was observed. This indicates that while the genetic backgrounds of the source populations were mostly retained, recombination introduced useful variation. Overall, the clear population structure and high diversity observed among these DHLs provide a strong genetic foundation for future maize improvement. These lines represent valuable resources for heterotic group formation, hybrid development, and recurrent selection schemes aimed at enhancing drought and low nitrogen tolerance in tropical maize.

## Introduction

Maize (*Zea mays L*.) is the most widely cultivated cereal globally, playing a vital role in food security, livestock feed, biofuel production, and industrial applications (Shiferaw et al., 2011; Vathana et al., 2019). However, maize production faces numerous challenges, including diseases, drought, low nitrogen availability, and *Striga* infestation. Among these, drought and low nitrogen stress are the most critical constraints affecting maize yields in sub-Saharan Africa. Developing climate-resilient maize varieties, particularly those with drought and low nitrogen tolerance, is essential for sustaining productivity in these regions. Over the years, the International Institute of Tropical Agriculture (IITA) and the International Maize and Wheat Improvement Center (CIMMYT) have conducted extensive research on breeding for drought and low nitrogen tolerance (Badu-Apraku et al., 2018; Talabi et al., 2017). However, understanding the genetic diversity of newly developed lines remains crucial for maximizing heterosis and selecting superior parental lines for breeding programs. Expanding the genetic base of maize germplasm for drought and low nitrogen tolerance is necessary to ensure the continuous development of stress-resilient varieties (Nelimor et al., 2020; Abu et al., 2021). Furthermore, efficient utilization of these genetic resources requires an in-depth understanding of their genetic diversity and population structure, which are key factors in optimizing genetic gains in maize improvement (Kumar et al., 2022; Mishra et al., 2023).

Genetic diversity studies provide insights into the relatedness of germplasm, allowing for the classification of inbred lines into subpopulations and heterotic groups (Billah et al., 2021; Gaytán-Pinzón et al., 2022). These studies can be conducted using morphological, biochemical, or molecular approaches. However, molecular markers provide a rapid and environment-independent method for assessing genetic relationships and variability (Cui et al., 2017; Osuman et al., 2020). Various marker systems, including simple sequence repeats (SSR), restriction fragment length polymorphism (RFLP), amplified fragment length polymorphism (AFLP), and single nucleotide polymorphisms (SNPs), have been used in maize genetic studies (Yu et al., 2011; Nisar & Hussain, 2022; Pecetti et al., 2023). Among these, SNP markers have gained prominence due to their abundance, stability, and high resolution in genome-wide analyses (Deschamps et al., 2012).

Diversity Arrays Technology sequencing (DArTseq) is a high-throughput genotyping method that combines next-generation sequencing with a genome complexity reduction step to efficiently discover and genotype thousands of genome-wide SNPs. DArTseq markers have proven to be a powerful tool for genetic diversity studies due to their ability to detect polymorphisms at a high density across the genome, making them highly useful for genomic selection and association studies (Sansaloni et al., 2011; Kilian et al., 2012). The application of DArTseq in maize breeding has facilitated precise population structure analysis and the identification of genetic variations associated with key agronomic traits (Thuo et al., 2020; Beyene et al., 2021).

Several studies have explored the genetic diversity of tropical maize germplasm, revealing high levels of genetic variation, which is crucial for breeding programs. For instance, Adu et al. (2019) reported significant genetic variation in West African maize inbred lines using SNP markers, highlighting the potential for developing climate-resilient hybrids. Similarly, Akande et al. (2021) examined the genetic structure of maize populations from Eastern and Southern Africa, demonstrating distinct subpopulation clustering, which is critical for heterotic group formation. Research on doubled haploid (DH) lines has also confirmed considerable genetic diversity, with DH lines exhibiting strong differentiation based on their source populations (Gaytán-Pinzón et al., 2022; Semalaiyappan et al., 2023). These findings underscore the importance of characterizing DH lines to optimize their use in hybrid development. DH lines offer significant advantages in genetic studies and hybrid breeding, as they are 100% homozygous and can be directly evaluated as parental lines for hybrid development, including combining ability studies

This study aims to determine the genetic diversity and population structure of newly developed maize DH lines derived from five drought and low nitrogen tolerant populations, some of which are widely cultivated in West and Central Africa.

## Material and methods

### Genetic materials and DH line development

A panel of 11 open-pollinated varieties (OPV) were evaluated prior under stress conditions and the five OPVs used for this study were selected based on their responses to the stresses. DH lines were extracted from the five diverse open pollinated (OPV) maize populations (Early thaï, Suwan 1 and Obatanpa sourced from Senegal, 87036(A) sourced from Ghana and TZE-WDT C4-STR from IITA, Nigeria. Early thaï, Suwan 1 and Obatanpa are varieties cultivated by farmers in West and Central Africa while 87036(A) and TZE-WDT C4-STR are improved populations. The DH lines were developed at the International Maize and Wheat Improvement Center (CIMMYT) facility in Kiboko, Kenya, following the protocol described by Chaikam *et al*. (2019). About 1000 plants per population were crossed with haploid inducer lines, resulting in a total of 5000 crosses. The putative haploid kernels of each plant were identified at the seed stage using the (R1-Navajo (R1-nj) gene, where expression of the anthocyanin pigmentation was only present in the endosperm while the diploid seeds expressed the pigmentation in both the endosperm and embryo (Chaikam *et al*., 2019). The haploid seeds were then germinated under controlled conditions to achieve a coleoptile length of 2 cm, and treated with colchicine for 8 hrs to double the chromosome number as described by Deimling et al. (1997). The coleoptiles were rinsed for 20 minutes after the treatment and were planted in trays in a greenhouse. The seedlings were moved to the field for seed production at the 4-5 leaf stage. During flowering, each plant was self-pollinated to generate the DH seeds. The 250 DH lines selected for this study were based on the number of seeds that were available for planting (Table 1).

**Table 1.**
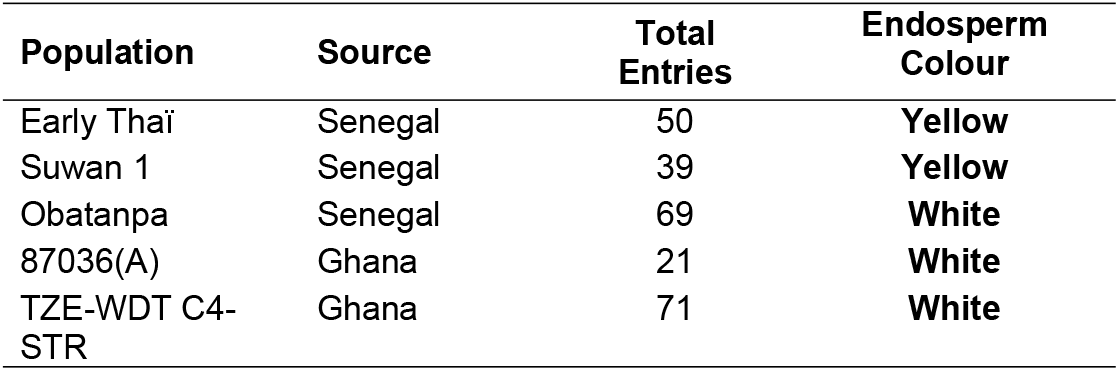
Description of the doubled haploid lines used in this study.

### Leaf sampling, DNA extraction and Genotyping using DArTseq technology

The 250 DH lines, comprising 161 with white endosperm and 89 with yellow endosperm, were sown in trays at a rate of two seeds per well. Leaf tissues were collected from seedlings 10 days after emergence, freeze-dried, and organized in 96-well plates before being sent to Intertek Australia for DNA extraction using the DArTseq protocol (Kilian et al., 2012; DArT Plant DNA Extraction Protocol. To assess DNA quality, electrophoresis was conducted on a 1% agarose gel, and degraded DNA samples were discarded. The DNA concentration was quantified using a NanoDrop 2000 spectrophotometer (Thermo Scientific, USA) and adjusted to a range of 50–100 ng/µL. The DNA was digested using the ApeKI restriction enzyme, following the Genotyping-by-Sequencing (GBS) protocol. Genotyping was performed using a mid-density platform, generating 3,305 SNPs. Sequencing was carried out on an Illumina HiSeq 2500 using a 96-plex GBS library (Elshire et al., 2011). The sequencing reads were aligned to the *Zea mays L*. reference genome, AGPv3 (B73 Ref-Gen v4 assembly) (Jiao et al., 2017). SNP calling and mapping were conducted using TASSEL 5.0 software. Initial filtering was applied to retain SNPs with a minor allele frequency (MAF) of at least 5% and a maximum missing data threshold of 20% (call rate ≥80%), while SNPs without known chromosomal locations were eliminated. This filtering process resulted in 2,135 SNPs retained for further analysis.

### Quality control and Filtering

To address missing genotype data and improve data quality for downstream genetic diversity analyses, imputation was performed using KDCompute ([KDCompute Platform](https://kdcompute.seqart.net/kdcompute/plugins)). An additional 10% of missing values were introduced into the dataset to evaluate different imputation methods. The imputation methods tested included Nonlinear Iterative Partial Least Squares (NIPALS), Probabilistic Principal Component Analysis (PPCA), Singular Value Decomposition (SVD), K-Nearest Neighbor (KNN), Expectation Maximization (EM), and MissForest. Each method’s effectiveness was assessed using the Simple Matching Coefficient (SMC), with MissForest yielding the highest SMC score. Consequently, the MissForest-imputed dataset was selected for genetic diversity analysis (Table S1).

### Data analysis

The Minor Allele Frequency (MAF), Polymorphic Information Content (PIC), pairwise Euclidean genetic distance (GD), kinship matrix, and markers (SNP) distribution to obtain the genotype and taxa summaries were computed using Tassel and PowerMarker software. The population structure was assessed based on admixture analysis in LEA package in R statistical software.

To further explore the genetic diversity among the DH lines, the data was subjected to the discriminant analysis of principal component (DAPC) that combines successive means of K and model selection to describe and infer clusters into populations of individuals that are genetically related (Jombart *et al*., 2010) in Adegenet package in R. All the analyses were performed in R statistical Computing Environment 4.2.1 (R Core Team, 2022).

## Results

### Summary statistics on the Population studied

The analysis of genetic diversity among the studied populations showed moderate variability across all parameters assessed. The MAF ranged narrowly between 0.28 and 0.30, with a mean value of 0.29, suggesting that both common and relatively rare alleles were represented within the populations (Table 2). Gene diversity values, which measure the probability that two randomly selected alleles are different, averaged 0.39, indicating a relatively high level of heterogeneity. Similarly, the Polymorphic Information Content (PIC), reflecting the informativeness of the markers used, ranged from 0.31 to 0.34, with an overall mean of 0.33.

**Table 2.** Summary genetic statistics of the source populations studied.

| Population | Sample Size | Minor Allele Frequency | Gene Diversity | Polymorphic Information Content |
| --- | --- | --- | --- | --- |
| Early Thai | 50 | 0.30 | 0.40 | 0.34 |
| Obatanpa | 69 | 0.30 | 0.40 | 0.33 |
| Suwan 1 | 37 | 0.29 | 0.38 | 0.32 |
| 87036(A) | 21 | 0.28 | 0.37 | 0.31 |
| TZE-WDT C4-STR | 71 | 0.29 | 0.38 | 0.33 |
| Means |  | 0.29 | 0.39 | 0.33 |

### Genetic grouping of the DH lines

The heatmap based on likelihood of identity by descent (IBD) yielded five clusters for 250 DH lines (Figure 1), and the DAPC identified 5 groups (Figure 2). Variables like kernel colour and line origin were utilized to create additional plots to better understand the relationship between inbred lines as depicted by the likelihood of IBD and DAPC. Although some lines were only partially grouped, distinct patterns between the origins of the lines and the endosperm colours were observed. The lines with the same endosperm colour tended to cluster together.

**Figure 1:**
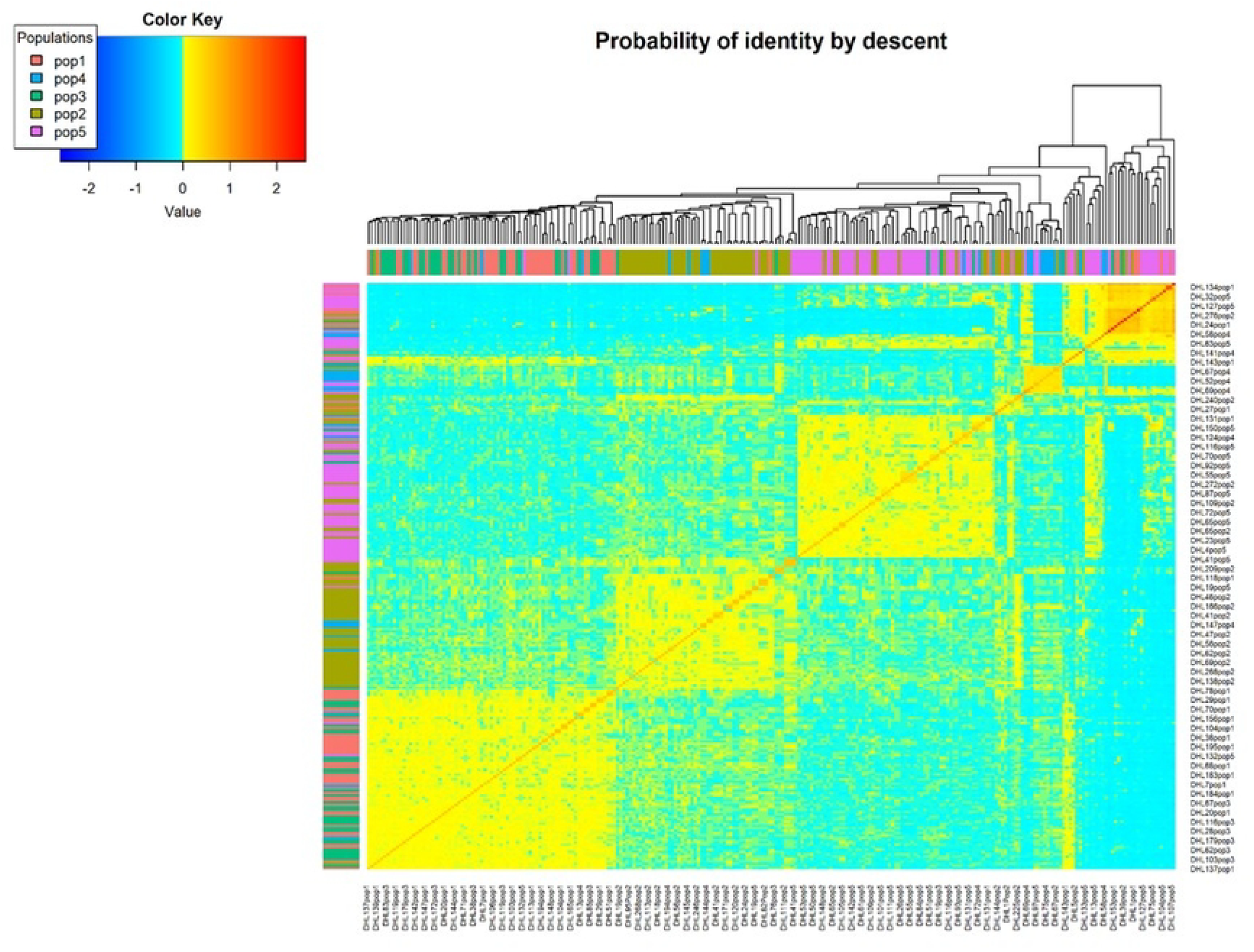
The heat map of SNPs among 250 DH lines based on genetic distances. The symmetric matrix representing the SNP rate was used for hierarchical clustering. The colours of the heat map correspond to the SNP rates, which are indicated in the legend at top left of Figure 1. A phylogenetic tree generated based on Euclidean distance is displayed above the heat map. The DH lines from the five populations are represented by five different colours. The distribution of the coefficient of ancestry is shown in the colour histogram, and the brighter the red hue, the more connected the individuals. The Bayesian Information Criterion (BIC) value with the lowest or most consistent value typically represents the best clustering. In this study, the lowest BIC value was 5, indicating that there were 5 distinct clusters as presented in Figure 2a. The BIC result showed that K = 5 was the best choice for grouping the lines into clusters. Five distinct clusters were also identified by the DAPC, and each cluster is indicated by a specific colour (Figure 2b).

**Figure 2:**
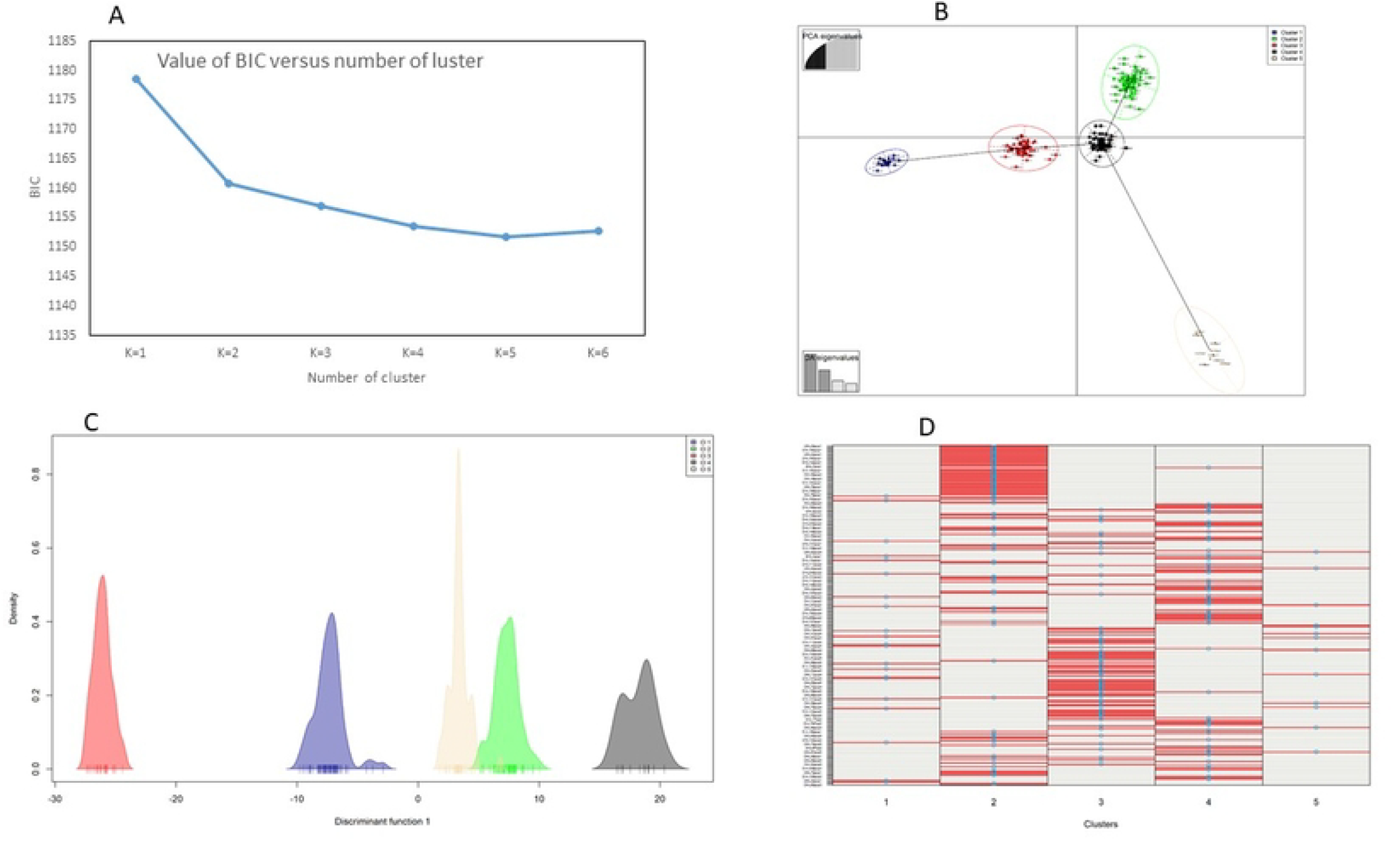
Genetic clustering of 250 DHLs by discriminant analysis of principal components (DAPC), (a) Bayesian information criterion (BIC); (b) Scatter plot from the DAPC. Cluster 1 (blue), Cluster 2 (green), cluster 3 (red) Cluster 4 (black) Cluster 5 (brown); (c) discriminant (d) assigned plot of the lines.

Four discriminant functions accounted for 77% of the variation in the overall dataset. Cluster 2, the largest cluster, consisted of 81 DH lines belonging to Populations 1 and 3, which were from the Senegal source. This was followed by Cluster 3 with 69 lines, primarily from the population 5 from the Ghana source. The lines from the Senegal source populations (Obatanpa, Early thaï, and SUWAN 1) comprised the majority of Cluster 4, which had 66 lines. The lowest number of DH lines were found in clusters 1 (21 lines belonging to population 5 from Ghana) and 5 (13 lines belonging to population 4 from Ghana). The first two linear discriminants are represented by the axes. The assigned plot as well as the discriminant function (Figure 2c) classified the lines in a similar pattern as the DAPC, confirming the five distinct clusters. In the assigned plot, the vertical bands depict the lines. The discriminant function also indicated 5 clusters. The number of lines in a group increases as the number of red bands and blue spots in a plot increases (Figure 2d).

The clustering analysis based on the DAPC components among the DH lines belonging to the white population indicated the presence of three discriminants (Figure 3a) and three clusters (Figure 3b). Cluster 1 comprised of 13 individuals belonging to 870436(A), that is population 4, while cluster 2 had 76 individuals, mostly belonging to population 5 (TZE-WDT C4-STR) and cluster 3 had 72 individuals belonging to population 2 (obatanpa) (Figure 3b). The grouping of the yellow DH lines based on the DAPC revealed two discriminants (figure 4a) two clusters (cluster 1 comprised of 16 lines, while cluster 2 comprised of 73 lines) (Figure 4b).

**Figure 3:**
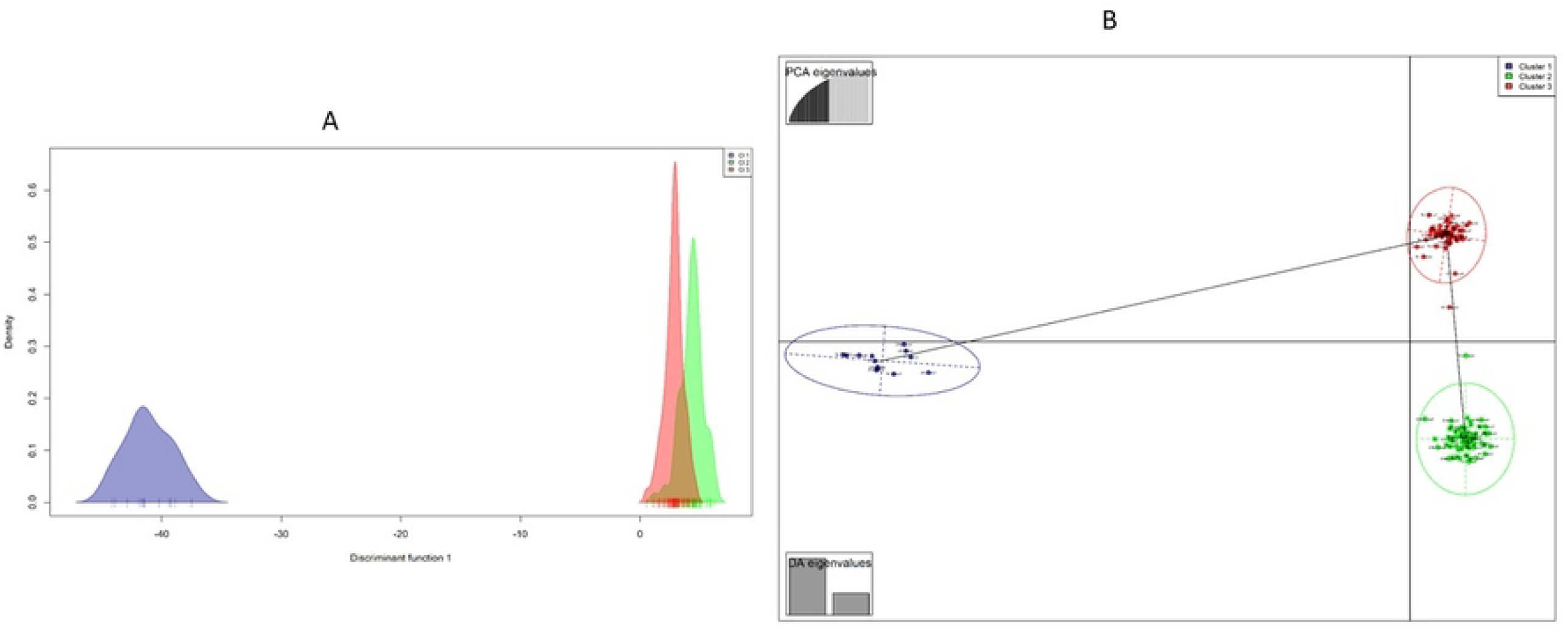
Genetic clustering of 161 white DHLs by discriminant analysis of principal components (DAPC), (a) discriminant function (b) Scatter plot from the DAPC. Cluster 1 (blue), Cluster 2 (green), cluster 3 (red).

**Figure 4:**
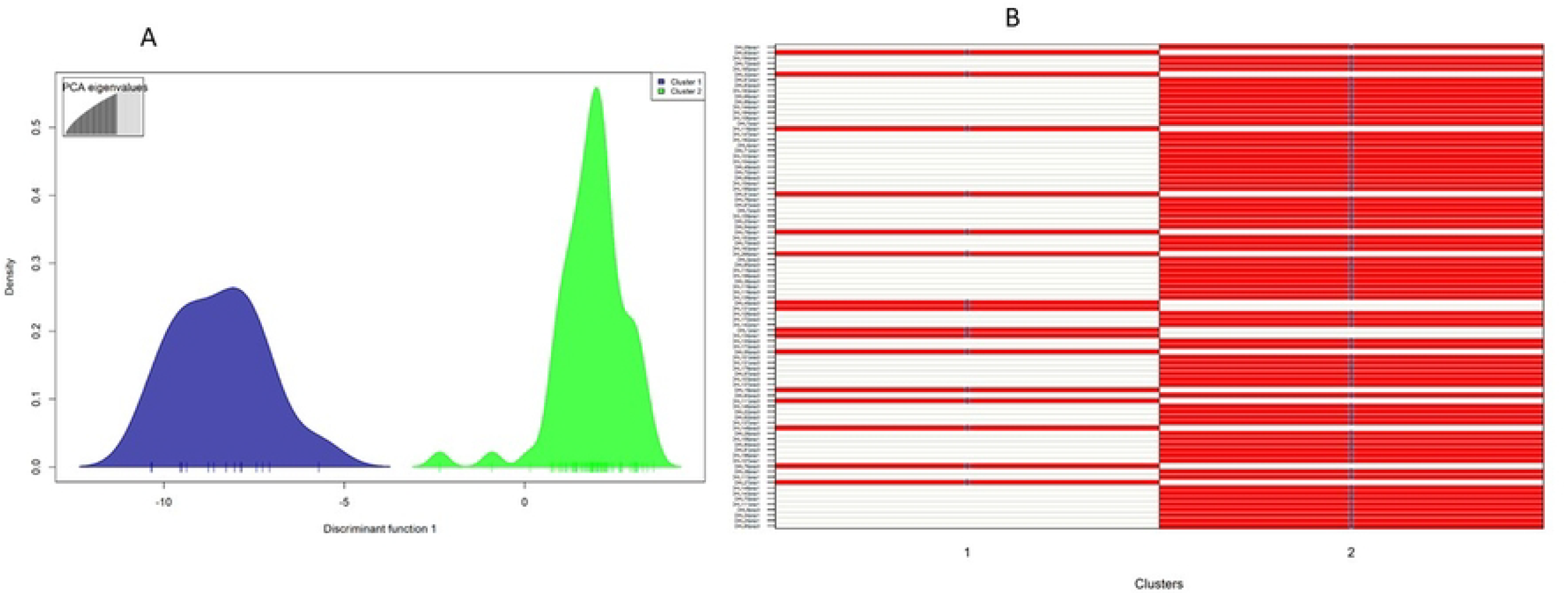
Genetic clustering of 89 yellow DHLs by discriminant analysis of principal components (DAPC), (a) discriminant function, Cluster 1 (blue), Cluster 2 (green) (b) assign plot from the DAPC.

The admixture analysis revealed the existence of five genetic sub-populations (Figure 5a). With the admixture model, when the membership probability proportion is greater than 0.5, the line is categorised and assigned to a group, whereas lines with a membership probability less than 0.5 were treated as admixed (Kumar *et al*., 2022). Sub-population 1 consisted of 7.6% of the DH lines largely from population 5 which is from Ghana. Sub-population 2 had 32% of the DH lines originating from population 1 and population 3 all with yellow endosperm colour and from Senegal. Sub-population 3 clustered 28.4% of the DH lines dominated by lines from population 5. Sub-population 4 grouped 22.4% of the DH lines from the Senegalese collection predominantly from population 2. Sub-population 5 contained 5.2% of the DH lines mostly from population 4. DH lines in admixture represent 3.9% of the total lines and consisted of lines developed from both Ghana and Senegal source populations (figure 5b).

**Figure 5:**
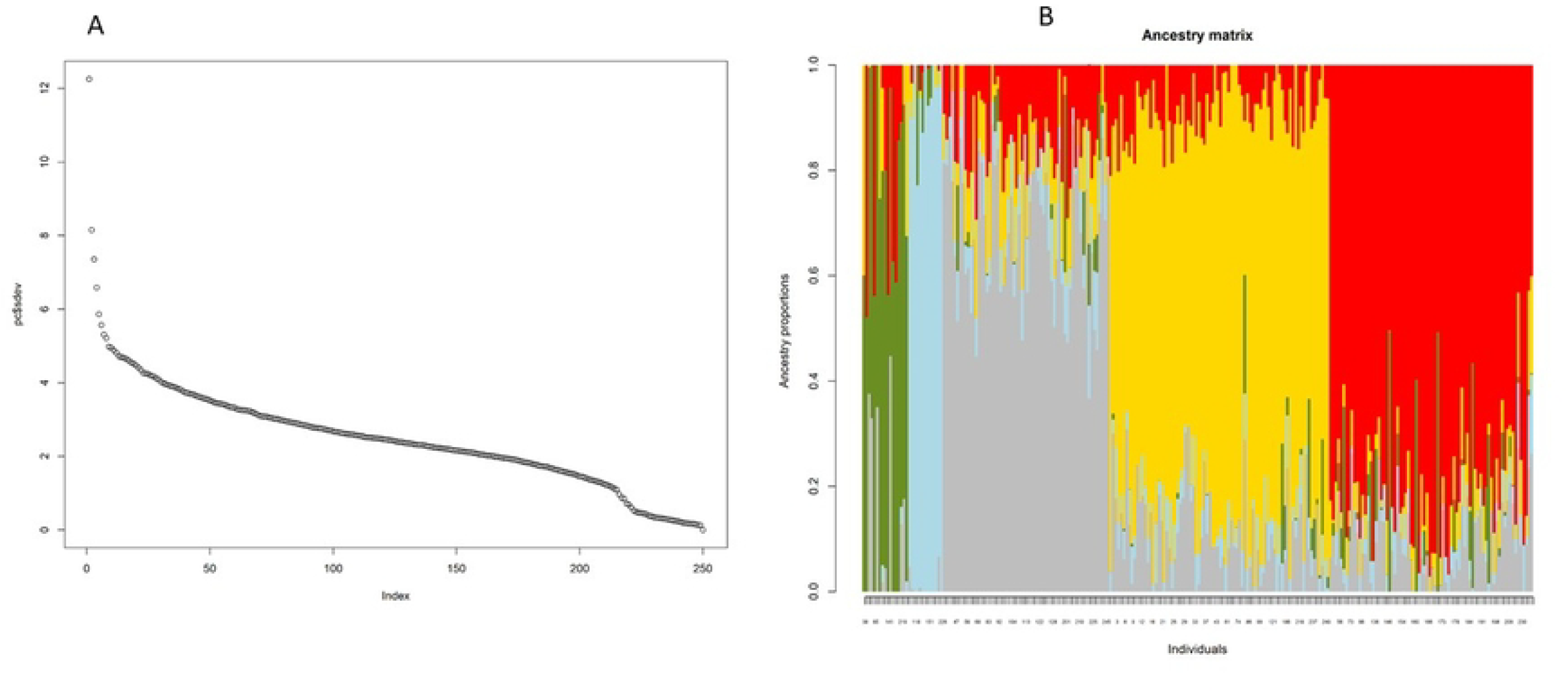
Population structure of 250 doubled haploid lines based on 3135 polymorphic at K = 5 SNP. (a) PC parameter for admixture, (b) admixture. Each doubled haploid line is represented by a vertical line that is partitioned into K-coloured segments. The colour green represents sub-group 1, blue corresponds to the sub-group 2, ash represents subgroup 3, yellow is sub-group 4 in, and red is sub-group 5.

### Molecular Analysis of variance among studied population

The Analysis of Molecular Variance (AMOVA) indicated that the majority of genetic variation (96%) was found within populations, while only 4% of the total variation occurred among populations. The estimated variance components were 656.236 within populations and 29.748 among populations, resulting in a total variance of 685.984 (Table 3). The PhiPT value, a measure of genetic differentiation among populations, was 0.043 and was statistically significant (P = 0.000), suggesting a low but significant genetic differentiation among the populations. The relatively higher variation within populations suggests substantial intra-population diversity, which is advantageous for breeding programs aiming to select diverse individuals (Table 3). The PhiPT value of 0.072 further supports the presence of some level of population structure, albeit weak.

**Table 3.**
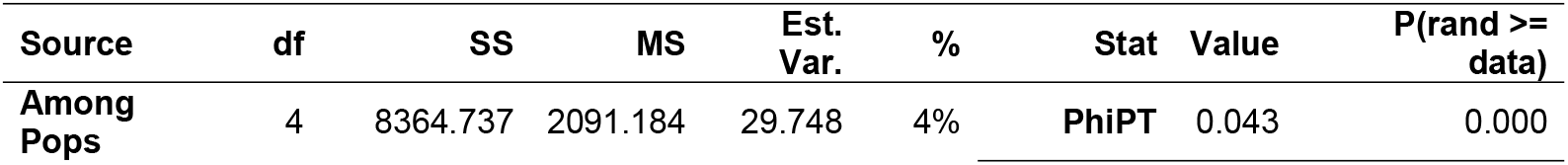
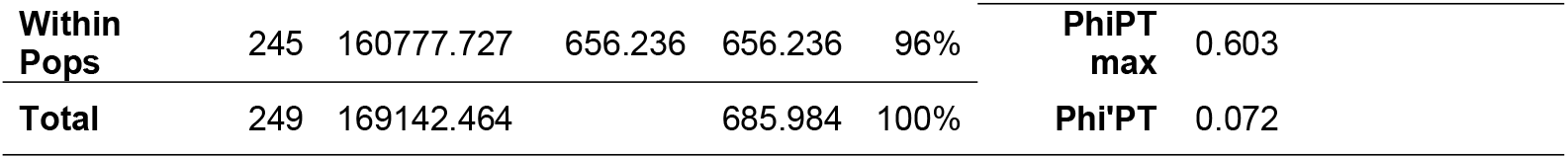
Analysis of molecular variance within and among the 250 populations with 3305 SNP markers.

| Source | df | SS | MS | Est. Var. | % | Stat | Value | P(rand >= data) |
| --- | --- | --- | --- | --- | --- | --- | --- | --- |
| Among Pops | 4 | 8364.737 | 2091.184 | 29.748 | 4% | PhiPT | 0.043 | 0.000 |

|  |  |  |  |  |  |  |  |
| --- | --- | --- | --- | --- | --- | --- | --- |
| **Within**<br>**Pops** | 245 | 160777.727 | 656.236 | 656.236 | 96% | **PhiPT**<br>**max** | 0.603 |
| **Total** | 249 | 169142.464 |  | 685.984 | 100% | **Phi'PT** | 0.072 |

## Discussion

Molecular markers have become reliable for identifying the genetic structure of germplasm (Adu et al., 2019). In this study, the genetic diversity of 250 DH lines extracted from five populations (two yellow and three white populations) was assessed using 2135 SNPs. The findings of this study indicate that the SNP markers have a PIC that ranged from 0.09 to 0.49 with an average PIC value of 0.3. This finding is similar to that reported (average PIC value of 0.354) by Gaytán-Pinzón et al. (2022) but is higher than average values (0.17-0.29) reported by Silva et al. (2020), Adu et al. (2019), Osuman et al. (2020), and Abu et al. (2021). These studies reported that the SNPs were sufficiently polymorphic for further genetic research. The relatively higher mean PIC value obtained indicated that the SNPs chosen for this study were sufficiently polymorphic and discriminatory for studying the inherent genetic diversity and capturing most of the genetic information among the DH lines. According to Gaytán-Pinzón *et al*. (2022), higher heterosis can occur when genetically distant inbred lines are crossed. Hence, future crosses for hybrid development should target lines that have the same endosperm colour but belong to different clusters to maximize the heterosis.

Information on genetic distance primarily reveals the diversity of the genes in haploid markers and provides information on the relatedness of individuals within and between populations (Luo *et al*., 2019; Osuman *et al*., 2020). The average GD of 0.39 measured in this study is comparable to previous values (0.30 - 0.39) in other maize inbred lines (Adu *et al*., 2019; Belalia et al., 2019; Obeng-Bio *et al*., 2020; Abu *et al*., 2021). From this finding, it can be inferred that the DH lines exhibited high genetic variability. This finding demonstrates that most of the DH lines in are distinctive and each is considered a viable candidate for contributing novel alleles to breeding programs, or via intercross to generate hybrids (Sodedji et al., 2021; Gaytán-Pinzón et al., 2022).

The SNPs had MAF that ranged from 0.05 to 0.29. The MAF between 0.05 and 0.25 obtained by approximately 41% of the 2135 SNPs for the 250 DH lines is lower than the MAF of 46% reported by Osuman et al. (2020) with 9684 SNPs employed on 162 lines. The value was however higher than 37% reported by Semagn et al. (2012) in 450 CIMMYT lines using 1065 SNPs. The DH lines are homozygotes and can be used as genetic resource in genetic studies and association mapping that require uniformity of the material.

Relative kinship which varies from 0 to 1.0 has been used largely to determine the similarity or dissimilarity of pairwise genetic relations (Cui *et al*., 2017). When the kinship value is close to 0 it means that the lines are unrelated (Wegary *et al*., 2019). In this study, 86% of the DH lines showed a pairwise mean relative kinship value of or close to 0. The result suggested low relatedness of the lines and the presence of little or no redundancy in the genomic composition of the DH lines, thus indicating large genetic diversity among the lines.

The hierarchical grouping and the SNP variations between the DHLs were used to create the heat map. Five groups were found in the heat map plot, which represented the likelihood of identification by descent. Generally, the lines from the same population were classified into the same group. Lines from population 1 and population 3, both from Senegal, clustered together. Lines from Population 2, from Senegal clustered with white lines derived from population 4 from Ghana. Population 4 has some pedigree relatedness with the 87036A, developed from a mid-altitude adapted line. Most of the source populations from which the DH lines were extracted were originally from the International Institute of Tropical Agriculture (IITA) hence they are expected to exhibit some relationship. Furthermore, it is noteworthy that endosperm colour also influenced the grouping of the lines.

The allocation of lines into heterotic groups is facilitated by the population structure analysis and the genetic distance. Individual descendants of the lines are used for this (Lawson and Falush, 2012). The result of the DAPC is consistent with the model-based population structure utilizing the BIC. Five potential sub-groups, or k = 5 was observed. The lines were grouped based on their similarity in terms of endosperm colour and origin. Most lines from the same population, with the same endosperm colour and parentage, tended to cluster into the same group. Nevertheless, some of the inbred lines from the same source population were not clustered together. This could be explained by the possible change in their genetic make-up. The outcomes demonstrated how well the markers classified the lines into different groups.

However, in this study there were discrepancies in the distribution of the lines into sub-groups in the population structure analysis and the DAPC analysis results. In contrast to the structural analysis, where many lines were considered as admixed individuals, the DAPC placed each DH line into a specific group. Similar findings have been reported by Sodedji et al. (2021), Campoy et al. (2016), Diouf et al. (2021), and Ketema et al. (2020) demonstrating the efficiency and precision of the DAPC. The five groups identified in this study are potential sources of beneficial alleles. The DArT-based SNP-derived markers were valuable in detecting differences among the lines, supporting findings of several authors (Ketema *et al*., 2020; Nyombayire *et al*., 2016); Obeng-Bio *et al*., 2020; Osuman *et al*., 2020; Wenzl *et al*., 2004). The DAPC classified the white DH lines into three different clusters and the yellow DH lines into two clusters. The clustering was efficient as the lines belonging to the same population were classified into the same cluster with a few exceptions. This indicate that the choice of the lines for hybrid development could be done by selecting lines from the same endosperm colour but from different clusters to maximize the heterosis.

The overall Fst of 0.37 reported by Mathiang et al. (2022) among sub-population in their genetic diversity and population structure study, was comparable to the Fst value, 0.2, obtained in this study. This could be explained by the differentiation due to the genetic structure of the populations used in this study. In breeding programs involving these set of DH lines, the ancestry should be considered in developing hybrids. The results of the diversity investigation showed that the lines are sufficiently diverse to be used in the association mapping of genes and candidates for breeding programs. The lines are also useful sources of favourable genetic resources for the increase of maize production in West and Central Africa.

## Conclusion

High genetic diversity and population structure were observed within the DH lines. Five sub-groups and five clusters were obtained in this study by both by the admixture and the DACP. The intra population DAPC revealed three clusters for the white doubled haploid lines and two clusters for the yellow doubled haploids lines. Each sub-group identified in this study can be regarded as having enough genetic diversity to serve as a future source for the generation of hybrids with each cluster as a putative heterotic group. The genotypic information and findings from this study are suitable for other genetic investigations, such as genome-wide association studies, and would be highly helpful for parental selection in new breeding programs.

## Abbreviation list

IITA: International Institute of Tropical Agriculture
CIMMYT: International Maize and Wheat Improvement Center
DArTseq: Diversity Arrays Technology sequencing
SNP: Single Nucleotide Polymorphisms
SSR: Simple Sequence Repeats
RFLP: Restriction Fragment Length Polymorphism
AFLP: Amplified Fragment Length Polymorphism DH: Doubled Haploid
MAF: Minor Allele Frequency
PIC: Polymorphic Information Content
GD: pairwise Euclidean genetic distance
NIPALS: Nonlinear Iterative Partial Least Squares
PPCA: Probabilistic Principal Component Analysis
SVD: Singular Value Decomposition
KNN: K-Nearest Neighbor
EM: Expectation Maximization
SMC: Simple Matching Coefficient
DAPC: Discriminant Analysis of Principal Component
IBD: identity by descent
BIC: The Bayesian Information Criterion

## Author contributions

**Georgina Lala Ehemba:** Conceptualization, Data curation, Formal analysis, Investigation, Methodology, Sofware, Writing - original draft, Writing - review & editing. **Pangirayi Tongoona:** Supervision, Writing - review & editing. **Pearl Abu:** Conceptualization, Validation, Writing - review & editing. **Bishwanath Das:** Supervision, Validation, Writing - review & editing. **Eric Yirenkyi Danquah:** Supervision, Writing - review & editing. **Beatrice Elohor Ifie:** Conceptualization, Methodology, Supervision, Writing - review & editing. **Mathieu Anatole Tele Ayenan:** Conceptualization, Data curation, Formal analysis, Writing - review & editing. **Priscilla Ribeiro:** Methodology, Writing - review & editing. **Manilal William:** Data curation, Formal analysis, Visualization. **Emmanuel Amponsah Adjei:** Conceptualization, Data curation, Formal analysis, Writing - review & editing. **Ana Luisa Garcia-Oliveira:** Funding Acquisition, Project administration, Resources, Validation.

## Acknowledgments

The authors send their appreciation to the team of Intertek Australia for the genotyping, to the team at CIMMYT Kiboko in Kenya for the doubled haploid lines development used in this work and to GIZ for funding this work through EiB CIMMYT (GIZ 17.7860.4-001.00). A special acknowledgement to the European Union through the Partnership for Training Scientists in Crop Improvement for Food Security in Africa (SCIFSA) for the scholarship offered. We are grateful to the University of Ghana via WACCI for hosting us and supporting part of this work.

